# High early embryo mortality and low hatching success observed in Aldabra giant tortoise populations

**DOI:** 10.1101/2025.05.07.652633

**Authors:** Alessia M. Lavigne, Richard Baxter, Eric Blais, Robert W. Bullock, Mark Brown, Christina Marques, Angelin B. Sanders, Nirmal Shah, Chris Tagg, Elisabeth Wareing, James Wareing, Nicola Hemmings

**Affiliations:** The University of Sheffield; Island Biodiversity Conservation Centre, University of Seychelles & Indian Ocean Tortoise Alliance; North Island Company Limited; Nature Seychelles; Save Our Seas Foundation; School of Life Sciences, University of KwaZulu-Natal & Cousine Island Company Limited; Cousine Island Company Limited; Island Conservation Society; University of Sheffield

## Abstract

In long-lived species, logistical constraints often limit monitoring to adult population censuses, potentially generating data that is biased towards older, more discernible individuals, and obscuring problems occurring at early life stages. Since adult populations of long-lived species can persist for decades despite reduced productivity, declines driven by reproductive failure may remain undetected until sudden population collapses occur. We present preliminary data on fertilisation success, early embryo survival, and hatching success, across one natural and five translocated populations of Aldabra giant tortoises (*Aldabrachelys gigantea*) in the Seychelles. Of 317 eggs from 24 clutches, only 16% successfully hatched. Most failed eggs (97%) were undeveloped, and using recently developed microscopic methods to assess egg fertility, we provide the first population-level comparisons of fertilisation and hatching outcomes for this species. Although sample sizes are limited, our results consistently indicate low and variable hatching success across populations, driven primarily by embryo mortality. Complete clutch failure was common (67%), and among clutches that did experience some degree of hatching success, early embryo mortality was still prevalent. Hatching rates were markedly reduced in translocated populations (0-26%) compared to the natural Aldabra Atoll population (46%) and consistently fell below historical hatching rate estimates (60-80%) reported ∼50 years ago. Although preliminary, these data provide the most accurate estimates of fertility available for Aldabra giant tortoises, as well as the first reproductive success data for any translocated population and the first for Aldabra in the last five decades. Our findings highlight the limitations of relying solely on adult census data for threat assessments in long-lived species. We argue that it is essential to incorporate productivity metrics into monitoring frameworks to improve population vulnerability predictions and inform effective conservation management.

## Introduction

Long-lived species play critical roles in ecosystem stability and functionality, often serving as key players in food web structures (Hazen et al., 2019; Kopf et al., 2024) and undertaking fundamental roles in environmental nutrient cycling and ecosystem engineering (Lovich et al., 2018; Albertson et al., 2024). They can provide insights into the evolutionary and physiological basis of longevity and age-related disease (Quesada et al., 2019; Omotoso et al., 2021), and many hold significant cultural and economic value (Kopf et al., 2024).

From a conservation perspective, understanding the population dynamics of long-lived species is crucial for informing estimations of population growth rates, future population sizes, and extinction risk (Briggs-Gonzalez et al., 2017; Jackson et al., 2020; Kopf et al., 2024). Life history traits (especially those related to pace of life and reproductive strategy) directly influence population dynamics and ultimately a species’ resilience in our current world of rapid global change (Capdevila et al., 2022). However, data on life-history traits of long-lived species (e.g., generation times, reproductive output, growth and maturation rates, and recruitment) are typically limited due to challenges in monitoring their populations over prolonged time periods (Briggs-Gonzalez et al., 2017; Edwards et al., 2019). Therefore, our ability to identify, manage, and predict potential population disturbances and to develop effective recovery strategies is often limited.

While modelling approaches can be used to predict future population trajectories of long-lived species, a comprehensive understanding of species life history and population demography is essential for informing these models (Benton et al., 2006). Population dynamics of long-lived species have traditionally been expected to be most sensitive to adult survival rate (Souchay et al. 2013), but in many cases, factors such as fecundity or juvenile survival are more influential. A study on the long-lived (30-86 years; Wiese & Willis, 2004) and endangered Asian elephant (*Elephas maximus*), using data collected over 40 years, showed that variation in age-specific survival and reproduction significantly impacted population growth rates (Jackson et al., 2020). Similarly, while adult survival was shown to be the most important factor influencing population growth in a 36-year study of American crocodiles (*Crocodylus acutus*), juvenile and subadult survival rates also had a significant influence (Briggs-Gonzalez et al., 2017). In a >15-year study of the imperilled bog turtle (*Glyptemys muhlenbergii*), increased survival rates at egg and juvenile stages were shown to buffer adult mortality and emigration rates, and in some populations, overall population decline was induced by reduced survival at multiple life stages, including the egg and juvenile stage (Knoerr et al., 2021). These findings demonstrate the complexity of population dynamics in long-lived species and the need for long-term data collection on factors such as population age structure and egg/juvenile survival to inform conservation and management strategies.

Species categorisation on the International Union for Conservation of Nature (IUCN) Red List of Threatened Species is arguably one of the most influential current drivers of conservation policy. The accuracy of the IUCN’s assessments relies on the quality of data available for each criterion, which is in turn dependent on monitoring effort and understanding of key life history traits. For species with limited data, assessments can still be made via “inference, suspicion, and projection”, provided assumptions are documented (IUCN Standards and Petitions Committee. 2024). Many IUCN Red List assessments are based on changes in population size, which is why risk assessments are often informed by census data (i.e., counts of mature individuals). However, even if census data are available for long-lived species, changes in reproductive output and population demography, which may undermine long-term population persistence, can be easily masked by the longevity of mature individuals. It is known that long-lived species can maintain seemingly stable populations despite low recruitment, and this can eventually lead to sharp population declines. A >20-year (1970s-1990s) multi-population study of long-lived hellbender salamanders (*Cryptobranchus alleganiensis*), for example, showed an average population decline of approximately 77%, starting in the 1980s and characterised by a sustained shift in age structure and density across all populations, with a disproportionate decline in young individuals and relative increase in larger, mature individuals over time (Wheeler et al., 2003). Proposed causes included lack of recruitment, reproductive failure, and/or low survival of eggs or young (Wheeler et al., 2003). Mark-recapture surveys corroborated the trend of rapid decline in hellbenders, with studied populations consisting nearly exclusively of large, older individuals with limited signs of reproduction (Burgmeier et al., 2011). In 2004, hellbenders were listed as ‘Near Threatened’ by the IUCN, but in 2022, guided by these and other studies (IUCN SSC Amphibian Specialist Group, 2022), the species was uplisted to ‘Vulnerable’ because of their estimated 30-50% decline over the past three generations. This demonstrates the importance of detailed demographic data for informing accurate threat status classification. More recently, the implementation of headstarting, where individuals are reared in captivity during the most vulnerable early life stages, before being released into the wild in an effort to increase survival and recruitment, has proved effective for rebuilding hellbender populations – specifically, the collection and captive rearing of wild eggs has resulted in up to seven times more individuals surviving to adulthood compared to natural reproduction and egg development (Kaunert et al., 2024).

The above examples collectively highlight the value of comprehensive, long-term demographic monitoring and the critical role that complementary early-life-stage survival and recruitment data play in sustainably maintaining populations of long-lived species. Inclusion of data on these life-history traits can improve the accuracy of species’ threat statuses, with important implications for the allocation of conservation and research funding, as well as for the design and implementation of recovery programs.

### >Case study: the Aldabra giant tortoise Aldabrachelys gigantea

The Aldabra giant tortoise is a well-known example of an exceptionally long-lived species, with an estimated lifespan of 100+ years (Quesada et al., 2019). Giant tortoises were once widespread across Indian Ocean islands, but the Aldabra giant tortoise now stands as the only surviving species in this region (Palkovacs et al. 2002; Austin et al. 2003), with Aldabra Atoll in Seychelles being home to the only naturally occurring (i.e., not introduced) population. Despite having faced near-extinction at the end of the 1800s due to anthropogenic pressures (Stoddart et al., 1979), the tortoise subpopulations of Aldabra (divided into the main islands of Picard, Polymnie [no tortoises], Malabar and Grande Terre) comprise a total of approximately 100,000 individuals, representing the greatest abundance of free-living giant tortoises on Earth (Bourn et al., 1999; Turnbull et al., 2015).

Our understanding of the historical distribution of tortoises in Seychelles is limited (Stoddart et al., 1979). It is known that from 1850 to 1990, there was a period of active translocations of the Aldabra giant tortoise to other Seychelles islands; however, the details of these movements are scantily documented up until the late 1970s (Gerlach et al., 2013). Although it is likely that all populations of the tortoises in Seychelles have Aldabran origin/ancestry (Çilingir et al., 2022), the historical details of parental lineage, age, abundance, and distribution over time of most free-ranging and captive tortoises are unknown (including individual/population-level details of when and where they hatched and records of individual transfers to other islands) The most recent publication of the Aldabra giant tortoise distribution and abundance across the Seychelles took place over a decade ago, covering the period 1990-2012 (Gerlach et al., 2013).

The IUCN Red List currently categorises Aldabra giant tortoises as ‘Vulnerable’ based on their restricted distribution, since most individuals of this species are on Aldabra (Tortoise & Freshwater Turtle Specialist Group, 1996). Any major negative ecological events on Aldabra would inevitably have serious consequences for the overall persistence of the species (Samour et al., 1987). It was hoped that translocations to other islands would result in multiple viable populations (Samour et al., 1987), securing the species’ future while also making them more accessible for scientific study and as a pilot model for future translocations to other Indian Ocean islands. Recent extinctions of several large and giant tortoise species have disrupted island ecosystems, particularly with respect to seed dispersal and herbivory (Hansen et al., 2010). Since Aldabra giant tortoises are heavy grazers and significant biomass contributors, with the capacity to alter landscapes and influence ecosystem health and function (Coe et al.,1979, Lovich et al., 2018), conservation practitioners have been using them (and other species) to restore missing ecological functions (Hansen et al., 2010, Gerlach et al., 2013, Stark & Galetti, 2024). Consequently, it is in the best interest of all conservation entities managing Aldabra giant tortoise populations to better understand their population demography and the critical role that early-life-stage survival and recruitment play in population persistence of this long-lived species.

Most current knowledge on the population ecology, dynamics, and reproduction of the Aldabra giant tortoise comes from an intensive research period conducted on wild subpopulations on Aldabra in the 1970s-1980s (Gibson & Hamilton, 1984). Concerns regarding the long-term viability of the tortoises were noted (Bourn et al., 1999), leading to the establishment of a long-term monthly monitoring program in 1998, which continues today (Turnbull et al., 2015). This has led to valuable insights into the flaws of past census data and the species’ population dynamics, with the most recent estimations demonstrating current population stability (Turnbull et al., 2015). However, Turnbull et al. (2015) presented data on population trends over 15 years, which is a relatively short portion of this species’ generation time, and did not measure reproductive success, productivity, or age structure. Earlier attempts to measure early-life survival and productivity by a two-year study on Aldabra (Swingland & Coe, 1979) reported hatching rates of 60% (Malabar) and 80% (Grande Terre), and infertility (undeveloped eggs) rates of 10-20% at both locations. However, significant challenges in identifying tortoise nests, finding hatchlings/younger individuals, and determining egg fertility (especially undeveloped eggs) prevented the continuation or expansion of this type of monitoring program.

Detecting major shifts in Aldabra giant tortoise population dynamics – specifically in age structure and productivity – is challenging unless the adult population is severely impacted (Gibson & Hamilton, 1984; Bourn et al., 1999; Haverkamp et al., 2017). Data on factors influencing offspring production, early life survival, and recruitment, are temporally limited and spatially incomplete (Swingland & Coe, 1979; Gibson & Hamilton, 1984). Given that climate change has been suggested to have the greatest impact on juvenile survival in tortoises (Haverkamp et al., 2017), it is possible that low reproductive productivity – currently unmonitored in this species – may lead to future population declines that are undetectable via census monitoring in the short to medium term. Furthermore, previous findings on Aldabra may not be applicable to populations on other Seychelles islands that experience different environmental conditions (e.g., topography, soil type, vegetation, predation pressures).

Here, we provide preliminary records of the following key reproductive productivity indicators for Aldabra giant tortoise populations across six Seychelles islands (including Aldabra): hatching success, clutch size, and fertilisation rates. Our data provide the first information on fertilisation and hatching success for any wild Aldabra giant tortoise population over the last 50 years, and the first-ever comparison across translocated island populations in the Seychelles. We also provide the most accurate information on fertilisation rates in these populations, using novel methods to detect fertilisation failure and early embryo mortality in turtles and tortoises (Lavigne et al. 2024).

## Materials and methods

With the support of conservation partners, we obtained data on hatching success and egg fertility from as many clutches as possible in a 2-year period across six Seychelles islands. In 2022/2023, samples were collected from Aldabra: Picard Island (Seychelles Islands Foundation; 9.3910° S, 46.2139° E); North Island (North Island Company Limited; 4.3950°S, 55.2453°E); and D’Arros Island (Save Our Seas Foundation: D’Arros Research Centre (SOSF-DRC); 5.4180°S, 53.2962°E). In 2023/2024, samples were collected again from D’Arros Island, along with Cousin Island (Nature Seychelles; 4.3315°S, 55.6620°E); Cousine Island (Cousine Island Company Ltd; 4.3507° S, 55.6475° E); and Desroches Island (Island Conservation Society; 5.6952° S, 53.6583° E). Partners identified as many tortoise nests (n = 24) as possible during these breeding seasons, and at the end of incubation (∼6-8 months), they excavated nests, collected eggs that showed no sign of embryonic development for fertility analysis, and recorded all egg fates. Partners also counted the number of eggs in a nest during laying or post-incubation excavations to determine clutch size, and hatching success was calculated as the number of hatched eggs / total clutch size.

We chose an incubation time of ∼6-8 months to accommodate the variable time range of incubation between and within different populations, depending on environmental conditions. For example, on the Aldabra Atoll, reported incubation durations range from as little as 98 days (about 3 months) to 148 days (about 5 months; Swingland and Coe, 1979), whilst some partners and zoos (e.g., Paignton and Barcelona Zoo) report incubation times up to ∼8 months. Allowing an incubation period of ∼6–8 months accommodates for this uncertainty and reduces the risk of mistaking any viable eggs as failed.

Due to significant challenges with identifying active Aldabra giant tortoise nests, our sampling approach for undeveloped eggs differed between the 2022/2023 and 2023/2024 breeding seasons. In 2022/2023, we aimed to sample 1-2 undeveloped eggs per clutch at random, from as many clutches as possible. However, since nest location proved difficult for our partners, in 2023/2024, we opted to collect as many undeveloped eggs as possible from any clutches found.

We determined the fertilisation status of undeveloped eggs as described in Lavigne et al., (2024). Briefly, we used microscopy-based techniques to identify embryonic nuclei on the perivitelline membrane (PVM) of undeveloped eggs to confirm egg fertilisation. We considered eggs to be unfertilised if most of the PVM surrounding the yolk was retrieved, but no embryonic nuclei and minimal/no sperm were found consistently across all PVM sections.

If we couldn’t retrieve sufficient PVM to confidently determine fertility status (i.e., to clearly identify embryonic nuclei), we classified eggs as inconclusive.

### >Animal ethics and permits

This research was approved by the Seychelles Bureau of Standards to be carried out with Seychelles conservation organizations (Ref: A0157). Non-viable Aldabra giant tortoise eggs were received opportunistically from Seychelles-based collaborators and were authorized for export as per the agreement made with the Ministry of Agriculture, Climate Change and Environment, in accordance with Article 15 of the Convention of Biological Diversity. Additionally, permits were acquired under the Convention of International Trade in Endangered Species of Wild Fauna and Flora (CITES; Seychelles export permit #A1615, #A1622, #A1777; UK import permit #631966/03, #24GBIMPBRJYT2).

## Results

In the 2022/2023 breeding season, partners collected 18 undeveloped eggs from 15 different naturally occurring clutches across Aldabra (Picard), North Island, and D’Arros. In 2023/2024, partners collected 48 undeveloped eggs from 12 different clutches from D’Arros, Cousin, Desroches, and Cousine (total of 66 undeveloped eggs from 27 clutches). Clutch data (i.e., information on clutch size and egg fates) was collected for 24 of these clutches.

We found hatching success varied considerably across Aldabra giant tortoise populations (Table 1) but appeared to be generally low, based on the clutches our partners were able to find. On average, 67% (16/24) of monitored clutches suffered complete hatching failure, but success rates varied between islands, from 46% on a naturally wild subpopulation of Aldabra (Picard Island) to 0% on Cousine, Desroches, and North Island (all of which are translocated populations). Out of 24 monitored clutches (total of 317 eggs), only 16% of eggs successfully hatched (52/317) and 82% of (261/317) did not (4 eggs were unaccounted for). Most unhatched eggs were undeveloped (253 eggs), except for eight unhatched eggs across three clutches from Aldabra (Picard) that displayed visible signs of embryonic development. Therefore, undeveloped eggs accounted for 97% (253/261) of all failed Aldabra giant tortoise eggs.

**Table 1.** Clutch data from 24 different Aldabra giant tortoise clutches on a naturally occurring wild subpopulation of Aldabra Atoll (Picard Island) and five other Seychelles islands with translocated populations was used to calculate the number of clutches experiencing total clutch failure (i.e., 0% hatching success) as well as their average, range and median clutch size and hatching success. Data were collected from Aldabra and North Island during 2022/2023; Cousin, Desroches and Cousine from 2023/2024; and D’Arros samples were collected across both 2022/2023 and 2023/2024 nesting seasons.

| | Number of clutches | Mean clutch size (± standard error) | Number of total clutch failures | Mean % Hatching success (all eggs) | % Hatching success range (per clutch, min–max) | Median % hatching success (all eggs) |
| --- | --- | --- | --- | --- | --- | --- |
| Aldabra Atoll (Picard) | 7 | 10.3 (± 1.8) | 2 | 46 | 0–91 | 44 |
| Cousin | 3 | 13.0 (± 3.5) | 2 | 26 | 0–78 | 0 |
| Cousine | 3 | 16.3 (± 3.8) | 3 | 0 | 0–0 | 0 |
| D'Arros | 4 | 17.0 (± 2.5) | 2 | 11 | 0–36 | 3 |
| Desroches | 3 | 10.7 (± 2.3) | 3 | 0 | 0–0 | 0 |
| North Island | 4 | 14.3 (± 3.5) | 4 | 0 | 0–0 | 0 |
| **TOTAL** | **24** | **13.6 (± 0.5)** | **16** | **19** | **0–91** | **0** |

Across both breeding seasons, we determined the fertility status of 66 undeveloped eggs collected from 27 clutches across the six Seychelles islands (Figure 1). We found that fertilisation rates also varied across populations: all undeveloped egg samples we were able to conclusively examine from Aldabra (natural population), Cousin, and Desroches were fertilised, and most undeveloped eggs examined from Cousine and D’Arros were also fertilised, but we found no evidence of fertilisation in any conclusively examined undeveloped eggs from North Island. Therefore, in most populations, hatching failure in Aldabra giant tortoises is likely to be predominantly driven by embryo mortality, but fertilisation failure may pose a problem for some translocated populations (Figure 1 and Table 1).

**Figure 1.**
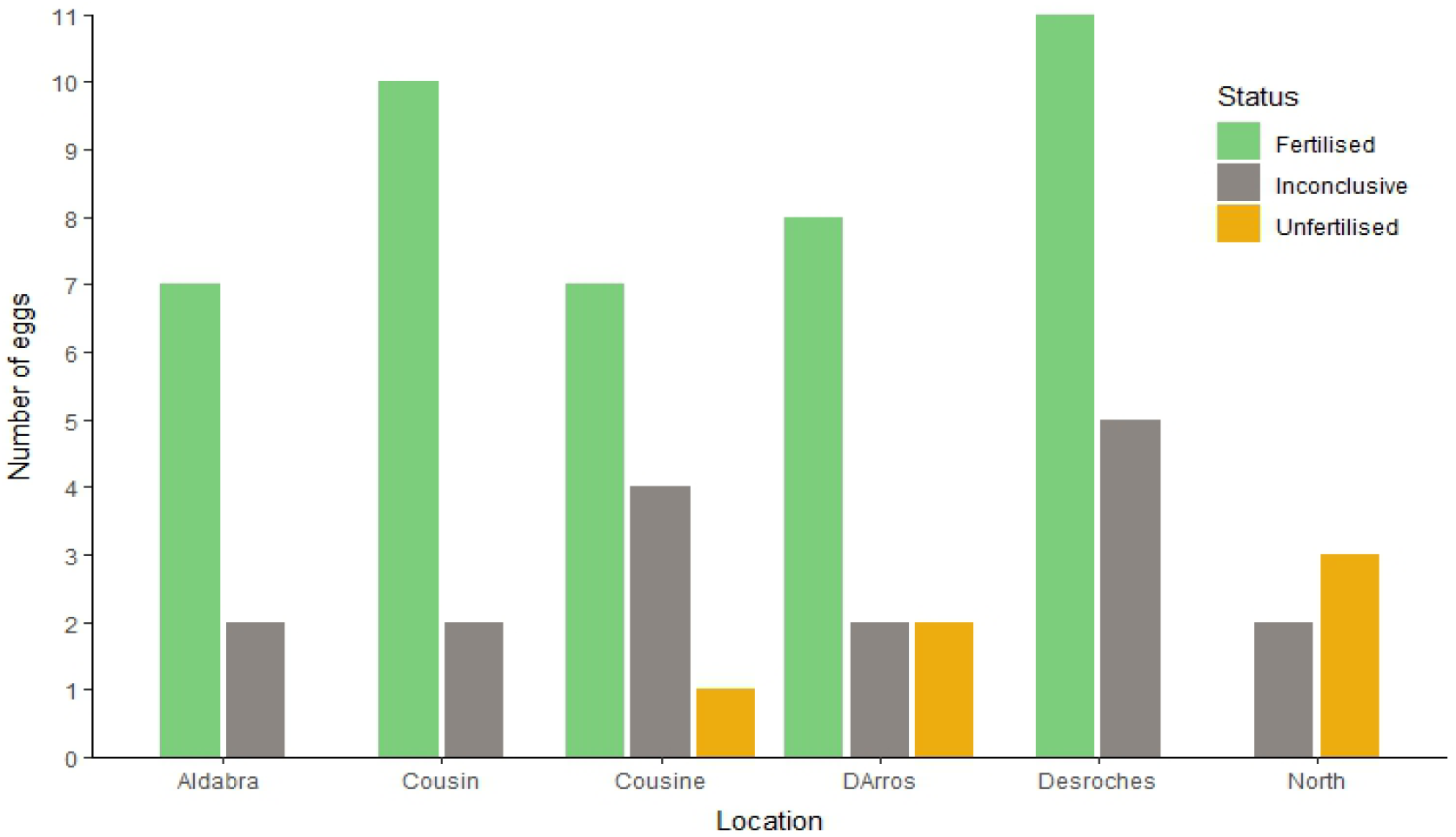
Fertility statuses of undeveloped eggs produced by Aldabra giant tortoises (Aldabrachelys gigantea, n= 66 eggs from 27 clutches) from six island locations in the Republic of Seychelles. Eggs were deemed inconclusive if there was insufficient material for further analysis. Samples were collected from Aldabra and North Island during 2022/2023; Cousin, Desroches and Cousine from 2023/2024; and D’Arros samples were collected across both 2022/2023 and 2023/2024 nesting seasons.

## Discussion

Functional declines in the populations of long-lived species, driven by low reproductive success and productivity, may be masked by apparently stable or even increasing (in the generational short-term) adult population sizes that persist due to long adult lifespans. Here, we present new data on early-life survival in the Aldabra giant tortoise, including fertility rates, early embryo survival, and hatching success across multiple populations in the Seychelles. Due to challenges with locating nests, our data are limited and should be considered preliminary. However, they are nonetheless suggestive of very low hatching success across multiple populations, particularly those translocated to the other Seychelles islands monitored in this study. We argue that more detailed monitoring of hatching success, fertilisation rates, and recruitment is needed to inform vulnerability assessments in this and other long-lived species.

Capitalising upon recent methodological developments (Lavigne et al. 2024), our study is the first to discriminate between fertilisation failure and early embryo death rates in Aldabra giant tortoises and compare these across populations ranging from a naturally occurring population on Aldabra to five other translocated populations on other Seychelles Islands. This novel approach reveals that early embryo death is likely the main barrier to hatching success, with fertilisation failure also presenting an issue in specific populations – particularly on North Island, where we found no evidence of fertilisation success in the four examined clutches. However, further investigation is warranted across multiple nesting seasons to fully understand these potential reproductive issues, since hatchlings, hatched eggs, and dead embryos have occasionally been found on North Island in the past (although not reliably recorded or monitored over time; AS pers. obs.). Multiple factors may contribute to both low fertilisation and embryo survival rates, and these need to be better understood through ongoing monitoring of fertilisation and hatching rates.

Although we sampled only a small number of eggs/clutches per population, relatively low hatching success was consistent across populations, with most clutches suffering high levels of early embryo death (Table 1; Figure 1). Determining a clear baseline of ‘normal’ hatching success for each island is challenging, as there is only a single study that has previously recorded hatching success levels on Aldabra (Malabar 60%, n = 213; Grande Terre 80%, n = 206), based on just 2 breeding seasons almost 50 years ago (Swingland & Coe, 1979). However, our hatching success data for Aldabra (Picard) (46%), Cousin (26%), D’Arros (11%), and Cousine, Desroches and North Island (all 0%) consistently fall well below range of 60-80% reported in this earlier study. Further investigations and larger sample sizes are required to determine the full extent of this variation and whether it depends on differences in terrestrial ecosystems or other (potentially anthropogenic) factors.

A previous baseline for fertilisation rates from Aldabra’s natural populations is even more elusive than that for hatching success, with only “a crude estimate of infertility” inferred from the percentage of clutches with undeveloped eggs only in Malabar (10%) and Grande Terre (20%) (Swingland & Coe, 1979). However, as new microscopic-based methods have demonstrated (Lavigne et al., 2024), it is inaccurate to assume that undeveloped tortoise eggs are unfertilised, drawing these previously reported infertility rates into question. We did not receive any undeveloped eggs from Malabar or Grande Terre to allow direct comparison with Swingland & Coe’s (1979) estimates, but our more accurate methods for determining fertility indicate that all undeveloped eggs from Aldabra (Picard) suffered early embryo death rather than fertilisation failure (Figure 1).

We can make a more direct comparison of clutch sizes of the Aldabra (Picard) population, since Swingland & Coe (1979) reported a mean clutch size of 19.2 eggs on Picard (n = 12 clutches across 2 years), whereas we found clutch size to be 10.3 eggs (n = 7 clutches in one year) in the same location (Table 1). Admittedly, both studies are temporally limited with relatively small sample sizes, but this is the only data currently available. While we have demonstrated the feasibility of obtaining more accurate baseline data on fertilisation rates, larger sample sizes across multiple breeding seasons are required to provide reliable estimates of clutch size and hatching success.

Monitoring population demography and life-history traits is becoming increasingly important amid rapid climate change. There is already evidence that the productivity of Aldabra giant tortoises may be impacted by environmental change: clutch size, egg weight, number of clutches per female, and rates of hatchling production and quality have all been shown to vary with rainfall, population density, and resource availability (Gibson & Hamilton, 1984; Swingland & Coe, 1979). Recent increases in drought frequency on Aldabra have also been posited to have a greater negative impact on juvenile survival/recruitment than adult survival (Haverkamp et al., 2017). It is therefore possible that climate change is creating suboptimal conditions for embryo development, thereby increasing embryo mortality (Figure 1). Increased climatic variability may also have contributed to shifts in life-history traits, such as clutch size. Our data suggest that, at least on Aldabra (Picard), clutch size may have both declined and become more variable compared to previous reports (Swingland & Coe, 1979). However, since productivity monitoring has been absent, it remains unclear how rapidly this change has occurred.

The absence of hatchlings/juvenile tortoises in population censuses may be due to their actual physical absence, or it could alternatively be explained by sampling bias, caused by difficulty in finding small/young tortoises. During Desroches’ first census in 2025, for example, even individuals weighing 9-15 kg were difficult to find, but it was later found that most newly found tortoises were in this size category (EW and JW pers. obs.). A similar sampling bias towards older, more conspicuous individuals is a well-known challenge during population censuses on other Seychelles islands (Gerlach et al., 2013). Additionally, significant portions of historical census and monitoring data across multiple islands are considered unreliable, including that of Aldabra and Frégate (Gerlach et al., 2013), the two islands with the largest population of Aldabra giant tortoises. Frégate Island – which was not included in this study – represents one of the most successful translocated populations, experiencing notable growth in recent years (Gerlach et al., 2013) from a small founder population of 40 individuals, to over 3500 free-roaming tortoises, thanks to habitat restoration, protection from humans, and additional reintroductions (R.B., pers. obs.). Despite the observed population growth on Frégate, monitoring of age structure, nesting activity, reproductive success, and overall productivity remains limited, and it remains unknown how previous and current trends in productivity have been shaped by pressures such as climate change. Complementing population census data with information on reproductive success would help provide a more comprehensive understanding of population dynamics, improving conservation decision-making and outcomes. One practical conservation solution would be to assess the potential for artificial incubation and headstarting of fertilised eggs from translocated populations of Aldabra giant tortoises on smaller islands like Cousin and Cousine, where hatching rates are currently low to non-existent in wild nests. If successful, headstarted individuals could boost recruitment to these and other populations.

Large, persistent declines and/or failures in reproductive success and recruitment have also been proposed as a potential future threat for critically endangered Galapagos tortoises (e.g., Western, *Chelonoidis porteri*, and Eastern, *C. donfaustoi, Santa Cruz Galapagos tortoises*; Blake et al., 2024), the only other extant giant tortoise taxa. Like the Aldabra giant tortoise, Galapagos tortoises are long-lived (Blake et al., 2024) and exhibit cryptic early-life-history stages. Their reproductive ecology (including hatching success, natal dispersal and recruitment rates) is poorly understood for similar technical and logistical reasons (Blake et al., 2024). However, in recent years, productivity and recruitment have been more intensively monitored. By measuring female condition, nest temperature, clutch characteristics, and survival, tracking individual movements using global positioning system telemetry and radiotelemetry, monitoring temperature and rainfall, and assessing primary productivity along elevation gradients, researchers have begun to elucidate how environmental variability influences female productivity and juvenile recruitment (Blake et al., 2024). The novel insights from this nine-year study will support the development of conservation strategies that consider tortoises’ unique reproductive ecology and slow life histories. Such valuable innovations in monitoring may be further enhanced by implementing methods for accurately determining fertilisation success and embryo survival rates, which have yet to be adopted for Galapagos tortoises. Such strategies are crucial for maximizing the resilience of critically endangered Galapagos tortoises in the face of projected rapid and dramatic shifts in land use, climate and invasive species in the Galapagos Islands; all of which are similar to what is expected in the Seychelles Islands, including Aldabra.

In summary, our work provides preliminary data that can provide a baseline for future studies of fertility and hatching success in Aldabra giant tortoises. It also highlights the need for finer-scale monitoring of reproductive success and productivity to complement current adult-focused population census studies (e.g., Turnbull et al. 2015), so that we can identify potential shifts in productivity that underpin longer-term changes in population demography. We require innovative approaches to facilitate this monitoring when species are long-lived and early life-history stages are cryptic. In the wake of new methods and technologies, future IUCN risk assessments should incorporate data on early reproductive success and productivity, especially for long-lived species such as the Aldabra giant tortoise. To facilitate this, however, appropriate funding and support for conservation and research monitoring must be a priority.

## Supporting information

The data supporting the findings of this study are openly available here

## Acknowledgments

This study was supported by a small grant 625 from Save Our Seas Foundation and small grant 237 from the Seychelles Conservation and Climate Adaptation Trust: Blue Grants Fund. We are grateful to the conservation teams and volunteers involved in sample collections and communications from Nature Seychelles on Cousin Island and the North Island Conservation Team (with special thanks to Mathilde Chloé Marie Le Gressus), DRC-SOSF team (specifically, Henriette Grimmel, Ellie Moulinie and Dillys Pouponeau), Seychelles Islands Foundation (with special thanks to the Aldabra Atoll team), Island Conservation Society: Desroches Island team, and the Cousine Island conservation team and its owner (Mr Keeley).

## Notes

### Competing Interest Statement

The authors have declared no competing interest.

### Summary of Updates

Although the results remain greatly unchanged, the manuscript's narrative and discussions underwent a large degree of rewriting following extensive reviewer feedback.

## References

Albertson, L. K., Sklar, L. S., Tumolo, B. B., Cross, W. F., Collins, S. F. & Woods, H. A. (2024). The Ghosts of Ecosystem Engineers: Legacy Effects of Biogenic Modifications. Functional Ecology, 38(1), 52–72.

Austin, J., Arnold, E. & Bour, R. (2003). Was There a Second Adaptive Radiation of Giant Tortoises in the Indian Ocean? Using Mitochondrial DNA to Investigate Speciation and Biogeography of Aldabrachelys (Reptilia, Testudinidae). Molecular Ecology, 12, 1415–1424.

Benton, T. G., Plaistow, S. J. & Coulson, T. N. (2006). Complex Population Dynamics and Complex Causation: Devils, Details and Demography. Proceedings of the Royal Society B: Biological Sciences, 273(1591), 1173–1181.

Blake, S., Cabrera, F., Cruz, S., Ellis-Soto, D., Yackulic, C. B., Bastille-Rousseau, G., Wikelski, M., Kuemmeth, F., Gibbs, J. P. & Deem, S. L. (2024). Environmental Variation Structures Reproduction and Recruitment in Long-Lived Mega-Herbivores: Galapagos Giant Tortoises. Ecological Monographs, 94(2), e1599.

Bourn, D., Gibson, C., Augeri, D., Wilson, C. J., Church, J. & Hay, S. I. (1999). The Rise and Fall of the Aldabran Giant Tortoise Population. Proceedings of the Royal Society of London. Series B: Biological Sciences, 266(1424), 1091–1100.

Briggs-Gonzalez, V., Bonenfant, C., Basille, M., Cherkiss, M., Beauchamp, J. & Mazzotti, F. (2017). Life Histories and Conservation of Long-Lived Reptiles, an Illustration with the American Crocodile (Crocodylus Acutus). Journal of Animal Ecology, 86(5), 1102–1113.

Burgmeier, N. G., Unger, S. D., Sutton, T. M. & Williams, R. N. (2011). Population Status of the Eastern Hellbender (Cryptobranchus Alleganiensis Alleganiensis) in Indiana. Journal of Herpetology, 45(2), 195–201.

Capdevila, P., Stott, I., Cant, J., Beger, M., Rowlands, G., Grace, M. & Salguero-Gómez, R. (2022). Life History Mediates the Trade-Offs among Different Components of Demographic Resilience. Ecology Letters, 25(6), 1566–1579.

Çilingir, F. G., A’Bear, L., Hansen, D., Davis, L. R., Bunbury, N., Ozgul, A., Croll, D. & Grossen, C. (2022). Chromosome-Level Genome Assembly for the Aldabra Giant Tortoise Enables Insights into the Genetic Health of a Threatened Population. GigaScience, 11, giac090.

Coe, M. J., Bourn, D. & Swingland, I. R. (1979). The Biomass, Production and Carrying Capacity of Giant Tortoises on Aldabra. Philosophical Transactions of the Royal Society of London. B, Biological Sciences, 286(1011), 163–176.

Edwards, J. E., Hiltz, E., Broell, F., Bushnell, P. G., Campana, S. E., Christiansen, J. S., Devine, B. M., Gallant, J. J., Hedges, K. J., MacNeil, M. A., McMeans, B. C., Nielsen, J., Præbel, K., Skomal, G. B., Steffensen, J. F., Walter, R. P., Watanabe, Y. Y., VanderZwaag, D. L. & Hussey, N. E. (2019). Advancing Research for the Management of Long-Lived Species: A Case Study on the Greenland Shark. Frontiers in Marine Science, 6. Retrieved from https://www.frontiersin.org/journals/marine-science/articles/10.3389/fmars.2019.00087

Gerlach, J., Rocamora, G., Gane, J., Jolliffe, K. & Vanherck, L. (2013). Giant Tortoise Distribution and Abundance in the Seychelles Islands: Past, Present, and Future. Chelonian Conservation and Biology, 12(1), 70–83.

Gibson, C. W. D. & Hamilton, J. (1984). Population Processes in a Large Herbivorous Reptile: The Giant Tortoise of Aldabra Atoll. Oecologia, 61(2), 230–240.

Hansen, D. M., Donlan, C. J., Griffiths, C. J. & Campbell, K. J. (2010). Ecological History and Latent Conservation Potential: Large and Giant Tortoises as a Model for Taxon Substitutions. Ecography, 33(2), 272–284.

Haverkamp, P. J., Shekeine, J., de Jong, R., Schaepman, M., Turnbull, L. A., Baxter, R., Hansen, D., Bunbury, N., Fleischer-Dogley, F. & Schaepman-Strub, G. (2017). Giant Tortoise Habitats under Increasing Drought Conditions on Aldabra Atoll—Ecological Indicators to Monitor Rainfall Anomalies and Related Vegetation Activity. Ecological Indicators, 80, 354– 362.

Hazen, E. L., Abrahms, B., Brodie, S., Carroll, G., Jacox, M. G., Savoca, M. S., Scales, K. L., Sydeman, W. J. & Bograd, S. J. (2019). Marine Top Predators as Climate and Ecosystem Sentinels. Frontiers in Ecology and the Environment, 17(10), 565–574.

IUCN SSC Amphibian Specialist Group. (2022). Cryptobranchus alleganiensis. The IUCN Red List of Threatened Species 2022: e.T59077A82473431. 10.2305/IUCN.UK.2022-2.RLTS.T59077A82473431.en. Accessed on 10 January 2025.

IUCN Standards and Petitions Committee. (2024). Guidelines for Using the IUCN Red List Categories and Criteria. Version 16. Prepared by the Standards and Petitions Committee. Downloadable from https://www.iucnredlist.org/documents/RedListGuidelines.pdf

Tortoise & Freshwater Turtle Specialist Group. (1996). Geochelone gigantea. The IUCN Red List of Threatened Species 1996: e.T9010A12949962. 10.2305/IUCN.UK.1996.RLTS.T9010A12949962.en. Accessed on 17 January 2025.

Jackson, J., Mar, K. U., Htut, W., Childs, D. Z. & Lummaa, V. (2020). Changes in Age-Structure over Four Decades Were a Key Determinant of Population Growth Rate in a Long-Lived Mammal. Journal of Animal Ecology, 89(10), 2268–2278.

Kaunert, M. D., Brown, R. K., Spear, S., Johantgen, P. B. & Popescu, V. D. (2024). Restoring Eastern Hellbender (Cryptobranchus a. Alleganiensis) Populations through Translocation of Headstarted Individuals. Population Ecology, 66(2), 93–107.

Knoerr, M., Tutterow, A., Graeter, G., Pittman, S. & Barrett, K. (2021). Population Models Reveal the Importance of Early Life-stages for Population Stability of an Imperiled Turtle Species. Animal Conservation, 25.

Kopf, R. K., Banks, S., Brent, L. J. N., Humphries, P., Jolly, C. J., Lee, P. C., Luiz, O. J., Nimmo, D. & Winemiller, K. O. (2024). Loss of Earth’s Old, Wise, and Large Animals. Science, 0(0), eado2705.

Lavigne, A., Bullock, R., Shah, N. J., Tagg, C., Zora, A. & Hemmings, N. (2025). Understanding Early Reproductive Failure in Turtles and Tortoises. Animal Conservation, 28(3), 353–364.

Lovich, J. E., Ennen, J. R., Agha, M. & Gibbons, J. W. (2018). Where Have All the Turtles Gone, and Why Does It Matter? Bioscience, 68(10), 771–781.

Omotoso, O., Gladyshev, V. N. & Zhou, X. (2021). Lifespan Extension in Long-Lived Vertebrates Rooted in Ecological Adaptation. Frontiers in Cell and Developmental Biology, 9. Retrieved from https://www.frontiersin.org/journals/cell-and-developmental-biology/articles/10.3389/fcell.2021.704966

Palkovacs, E., Gerlach, J. & Caccone, A. (2002). The Evolutionary Origin of Indian Ocean Tortoises (Dipsochelys). Molecular phylogenetics and evolution, 24, 216–27.

Quesada, V. et al. (2019). Giant Tortoise Genomes Provide Insights into Longevity and Age-Related Disease. Nature Ecology & Evolution, 3(1), 87–95.

Samour, H. J., Spratt, D. M. J., Hart, M. G., Savage, B. & Hawkey, C. M. (1987). A Survey of the Aldabra Giant Tortoise Population Introduced on Curieuse Island, Seychelles. Biological Conservation, 41(2), 147–158.

Souchay, G., Gauthier, G. & Pradel, R. (2013). Temporal Variation of Juvenile Survival in a Long-Lived Species: The Role of Parasites and Body Condition. Oecologia, 173(1), 151– 160.

Stark, G. & Galetti, M. (2024). Rewilding in Cold Blood: Restoring Functionality in Degraded Ecosystems Using Herbivorous Reptiles. Global Ecology and Conservation, 50, e02834.

Stoddart, D. R., Peake, J. F., Gordon, C. & Burleigh, R. (1979). Historical Records of Indian Ocean Giant Tortoise Populations. Philosophical Transactions of the Royal Society of London. B, Biological Sciences, 286(1011), 147–161.

Swingland, I. R. & Coe, M. J. (1979). The Natural Regulation of Giant Tortoise Populations on Aldabra Atoll: Recruitment. Philosophical Transactions of the Royal Society of London. Series B, Biological Sciences, 286(1011), 177–188.

Turnbull, L. A., Ozgul, A., Accouche, W., Baxter, R., ChongSeng, L., Currie, J. C., Doak, N., Hansen, D. M., Pistorius, P., Richards, H., van de Crommenacker, J., von Brandis, R., Fleischer-Dogley, F. & Bunbury, N. (2015). Persistence of Distinctive Morphotypes in the Native Range of the CITES-Listed Aldabra Giant Tortoise. Ecology and Evolution, 5(23), 5499–5508.

Wheeler, B. A., Prosen, E., Mathis, A. & Wilkinson, R. F. (2003). Population Declines of a Long-Lived Salamander: A 20+-Year Study of Hellbenders, Cryptobranchus Alleganiensis. Biological Conservation, 109(1), 151–156.

Wiese, R. J. & Willis, K. (2004). Calculation of Longevity and Life Expectancy in Captive Elephants. Zoo Biology, 23(4), 365–373.

